# Establishment of morphological atlas of *Caenorhabditis elegans* embryo with cellular resolution using deep-learning-based 4D segmentation

**DOI:** 10.1101/797688

**Authors:** Jianfeng Cao, Guoye Guan, Ming-Kin Wong, Lu-Yan Chan, Chao Tang, Zhongying Zhao, Hong Yan

## Abstract

Cell lineage consists of cell division timing, cell migration and cell fate, which are highly reproducible during the development of some nematode species, including *C. elegans*. Due to the lack of high spatiotemporal resolution of imaging technique and reliable shape-reconstruction algorithm, cell morphology have not been systematically characterized in depth over development for any metazoan. This significantly inhibits the study of space-related problems in developmental biology, including cell segregation, cell-cell contact and cell shape change over development. Here we develop an automated pipeline, CShaper, to help address these issues. By quantifying morphological parameters of densely packed cells in developing *C. elegans* emrbyo through segmentation of fluorescene-labelled membrance, we generate a time-lapse framework of cellular shape and migration for *C. elegans* embryos from 4-to 350-cell stage, including a full migration trajectory, morphological dynamics of 226 cells and 877 reproducible cell-cell contacts. In combination with automated cell tracing, cell-fate associated cell shape change becomes within reach. Our work provides a quantitative resource for *C. elegans* early development, which is expected to facilitate the research such as signaling transduction and cell biology of division.

## Introduction

Embryogenesis in metazoans is a spatio-temporal biological process formed by a series of multicellular structure evolution including proliferation and morphogenesis. As “eutely” *C. elegans* has invariant and reproducible cell lineage consisting of division timing, migration trajectory and fate specification for each cell^1^, it has been wildly used as an animal model for developmental biology research^2^. Thanks to advanced imaging equipment with single-cell resolution as well as automatic cell-tracing softwares^3–5^, a few researchers have made great effort to quantitatively reconstruct its developmental atlas in several dimensions of developmental properties, including cell division timing^6^, gene expression and morphogenesis^7,8^, cell-cell contact mapping and signaling^9,10^. Despite all this, little is known about cell morphology experimentally and systematically, due to lack of high-resolution cell membrane signaling marker and reliable imaging-based algorithm for cell segmentation, in particular for late stage which has hundreds of cells^11,12^. Cell morphology (e.g. cell shape, cell volume, cell-cell contact) is also a set of critical developmental properties and information for metazoan embryogenesis, which is tightly correlated to cell-cycle control^13–15^, spindle formation^16^, cell-fate symmetry breaking and differentiation^17,18^, intercellular signal transmission^10,12,19,20^, cytomechanics and morphogenesis^21–24^, etc.

Recent studies also emphasized the necessity of 3-dimensional cellular segmentation aside from the nucleus^25,26^. With large quantities of volumetric data involved in the embryogenesis of *C. elegans*, visual inspection on these images is time-consuming and error-prone. Computer-assisted analysis accommodates the difficulty in large-scale image segmentation, paving the way to efficient and accurate researches. Compared to manual annotation, automated image analysis has better objective quantification, consistency and reproducibility. Confocal microscopy is popularly used in 3D imaging, which allows optical section in tissue or even cells at different depth. Whereas large quantities of works have been proposed to segment nuclei or cells in 2D^27–30^, cell’s morphological features varies from those in 3D. Without information between slices, simply stacking 2D segmentations into 3D volume may yield misalignment in the depth direction. Some recent methods have addressed cell tracing based on nucleus information^31^, however, they can hardly characterize cell geometry information (e.g. cell volume, cell surface area, cell topology, cell-cell neighbour relationship). The performance of 3D cell segmentation suffers from three factors. First, unlike the nuclei, which are thick and separated ellipsoid components, cell membranes are thin planar structures, forming complicated networks by contacting with each other. Second, compared to plant tissue, highly dynamic cellular morphology in *C. elegans* limits the application of diverse techniques based on deformable model. Furthermore, as shown in Fig. S1**A**, laser attenuation makes the segmentation more challenging in deeper slices. Theoretically, large exposure times can improve the image quality. Poisonous laser rays, however, could harm or even kill cells in time-lapse imaging process.

In the last few decades, Several attempts have been made to leverage the segmentation performance on microscopy. Classical techniques are based on pre-defined model and image intensity features. Among these, active contours and level sets are two of the most compelling methods in segmenting microscopy. Active contour deals with segmentation as a energy minimization process where the image forces pull the contour toward the object boundary and internal forces resist the deformation. Different evolution equations, mediating the internal and external forces, are designed to control the deformation process precisely^32–35^. To diminish the difficulty in finding desirable forces representation, level set is used to embed the boundary curve as a real-valued solution of an equation, which makes it straightforward to follow topology changes, such as splits and holes. By using coupling constrains in level set evolution, Nath et al. proposed a computationally efficient method to segment hundreds of cells simultaneously^27^. Kiss et al. used level set to segment plant tissues at multiple scales, which reduced the error at blurry surface effectively^36^. Instead of processing multiple slices in 3D simultaneously, Sharp et al. described the possibility of inferring 3D shape features indirectly from 2D images in a statistical way^37^. In practice, however, the performance of level set is limited by considerable computational complexity and incomplete cell boundary. Methods driven by energy functions could be problematic due to the presence of artifacts and lack of strong edge information. Xing et al. provides a compressive review about classical cell segmentation techniques^38^. Data-dependent and parameterized pre-processing stage is always needed in these methods, otherwise the system would be exposed to under-or-over segmentation errors.

Recently, deep learning based methods provide a promising tool for recognition tasks, such as denoising^39–42^, tumor segmentation^43,44^ and image synthesis^45–48^. Compared to classical methods, convolutional neural network (CNN) shows remarkable improvement on biological image analysis by mining subtle texture and shape changes. Since U-net was proposed by Ronneberger et al.^49^, this kind of encoder-and-decoder structure has extensively promoted learning-based segmentations on medical images^50^. For fluorescent images, the ability of deep learning in discriminating and filtering useful information is further verified^48,51^. To mitigate the complexity in cellular networks, the segmentation is usually decomposed into multiple intermediate tasks, such as nucleus detection and membrane segmentation^**?**, 52,53^. 3D convolution attracts increasing attention because of its advantage in combining multi-directional information. However, 3D deep network relies heavily on computation resource and training data. Some works are proposed to alleviate the computational complexity in 3D^54–57^.

In this work, we propose a complete pipeline CShaper for analyzing cellular shapes in *C. elegans*. Deep learning is utilized to accommodate the complications associated in volumetric *C. elegans* embryo. First, instead of segmenting cells as a binary classification task directly, CShaper generates the discrete distance map from the membrane stack image with a distance regularized neural network DMapNet. Second, Delaunay triangulation is employed to construct a weighted graph based on the binary segmentation extracted from the DMapNet. Local minima are clustered as different seeds for watershed segmentations. Last, nucleus images are used to filter out hollow regions among cells. Automatic seeding procedure precludes substantial computations involved in most current works due to the over-segmentation problem. After adjusting position variations in wide-type emrbyos, we establish a spatio-temporal morphogenesis reference for *C. elegans* embryogenesis during 4-to 350-cell stage. Both individual evaluation and experimental verification on previous conclusions demonstrate the unprecedented performance of CShaper.

## Results

By measuring the consistency between segmentations of prevalent methods and manual annotations, CShaper outperforms regarding both accuracy and robustness. Besides, based on the segmentations, cell volume and cell surface area, which usually get involved in cell-cycle control and cell-fate specification^13–18^, were firstly investigated and found to have high reproducibility among individuals.

### Dataset

In *C. elegans* embryo, nucleus and membrane were stained in vivo with mCherry marker on nucleus and GFP marker on cell membrane, respectively. 4D (space + time) nucleus and membrane stacks from two channels were collected by Leisa SP8 confocal microscopy at 1.5-min interval. All images with a resolution 512 × 712 × 70 (voxel size × 0.09 × 0.43*µ*m) were resized into isotropic volume images with a resolution 205 × 285 × 134 (voxel size 0.22 × 0.22 × 0.22*µ*m) before analysis. Different wide-type embryos are used in different stages as listed in **Supplementary** Table. S1.

Manual annotations are needed to train the DMapNet and compare different methods. In plant tissue, cells are encased within cell walls that physically adhere to their neighbors, yielding better image quality and uniform size. In the animal embryo, however, irregular cells are separated by thin membrane, making it much challenging to segment each cell accurately. Besides, since only 2D slice can be shown on computer screen, full annotation of volumetric data is very tedious. Therefore, the gold standard is produced by semi-automatic software with the results revised by experts. Membrane stacks are pre-segmented by 3DMMS ^58^ first, and then checked by two experts with ITK-SNAP ^59^ slice-by-slice. Nucleus image is also imposed to prevent invalid gaps between cells. Please note that annotations are composed by cell-wise regions, which can be transformed into membrane mask with morphological operations. Most annotations have less than 100 cells to prevent annotation quality deteriorating with the image quality and segmentation error of 3DMMS. Although DMapNet is trained on frames within 150-cell stage, experiments show that it is able to process the embryo in 350-cell stage with high quality owning to the multi-scale input features. All datasets companied with annotations or segmentations will be publicly available.

### Comparison with other method

Here a through comparison among CShaper and other sate-of-the-art methods is discussed. To be a fair comparison, watershed algorithm is also used as a postprocessing procedure in 3D U-net and FusionNet where only binary membrane segmentation is available. However, different from CShaper, the seeds are collected from the nucleus locations generated by AceTree, which theoretically produces more accurate estimation on real nucleus images. In order to compare the CShaper without distance learning strategy, the CShaper was changed to a naive binary segmentation by replacing the last layer of DMapNet (see Methods) with two channel filters, while other parameters kept the same as that in CShaper.

Experimental results are reported in Fig. 1. It shows that CShaper achieves much higher accuracy on three different wide types at various time points (Fig. 1**A**), which validates the accuracy and generality of our approach. Such improvement benefits from: 1). CShaper allows segmentation on weak boundaries because of the distance constrained learning strategy (**Supplementary** Fig. S1); 2). In RACE and FusionNet, segmentations are implemented slice-by-slice. Although slices are combined into stack statistically during the post-processing stage, few inter-slice information, if any, in raw images are utilized to establish a promising result; 3). In contrast to naive binary segmentation methods, DMapNet reaches a compromise at the membrane boundary by constructing a relatively smooth transition between the membrane and the background. This encourages the network to learn more morphological features around the membrane.

**Figure 1:**
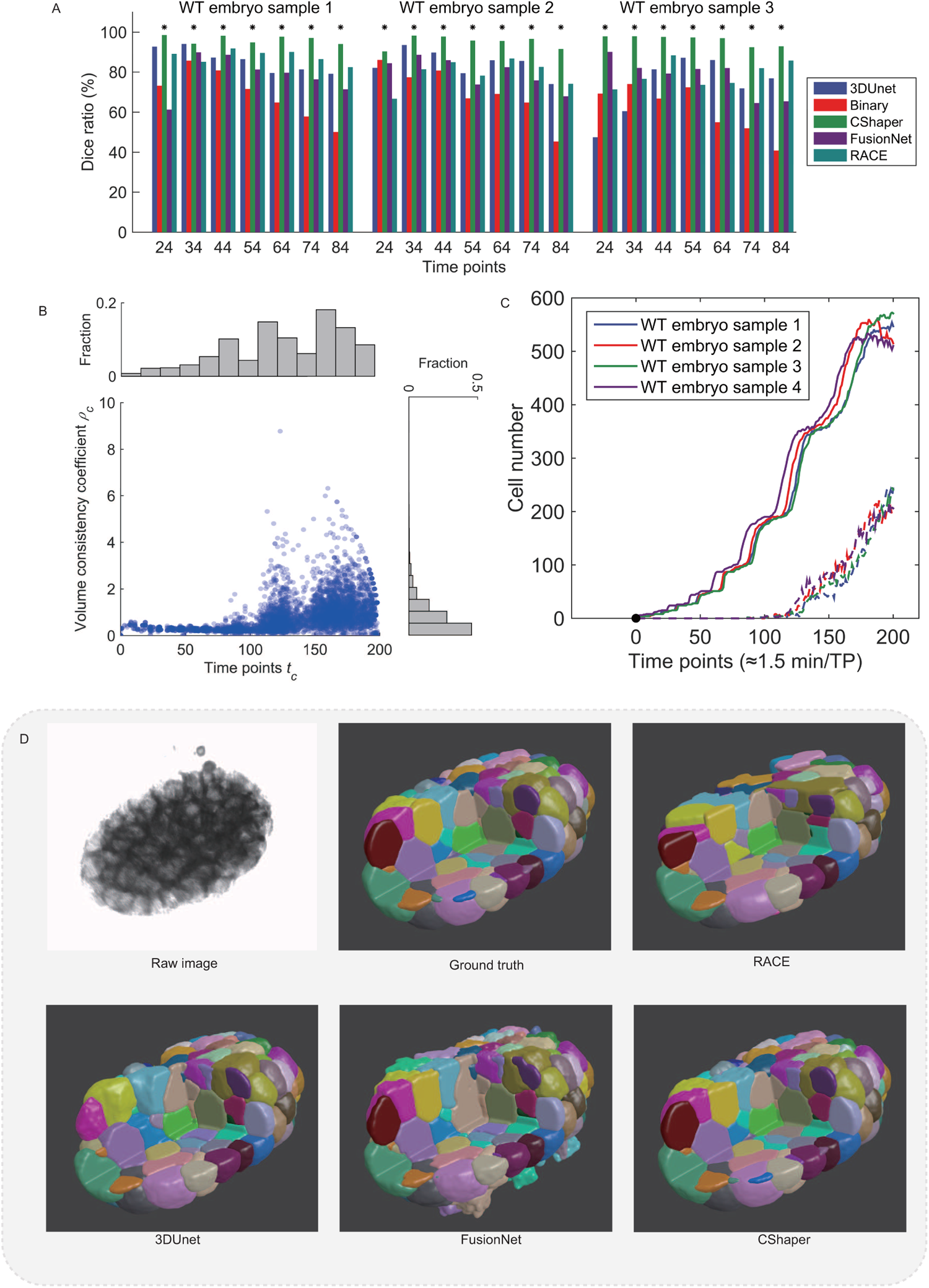
Analysis on the segmentation results. (**A**) Comparison on the Dice ratio of the segmentations from 3D U-Net, FusionNet, RACE, CShaper and naive binary. All values are calculated based on seven manual annotations of three wide types, respectively. (**C**) The distribution of cell volume consistency *ρ*_*c*_ with respect to time *t*_*c*_ in four embryos. (**D**) The distribution of lost nucleus with respect to time in four embryos. (**D**) Comparison of the segmentations from RACE, 3DUnet, FusionNet and CShaper.

As Fig. 1**D** shows, RACE and FusionNet suffer severe leakage at the button layers of the embryo, where the membrane signal is too weak to be discriminated because of laser attenuation. 3DUnet provides better feature extraction by applying convolution between slices. However, partial membrane, especially at the boundary of the embryo, is still too weak to be recognized, which deteriorates the lost cells in the periphery. Under the distance map constraint, CShaper learns to depict weak or lost membrane as annotated in the training data.

### Robustness on extensive datasets

To measure the performance of CShaper on more datasets, whose annotations are not available, we quantitatively measured the volume consistency, as well as the lost nucleus, of serial segmentations. CShaper segments each embryo independently, frame by frame, without capturing typical temporal patterns across time. Successive imaged cells, theoretically, should have temporally consistent volume, or with limited variance when considering biological dynamics. Therefore, we can examine the performance of CShaper on extensive time-lapse stacks by analyzing series information. Besides the membrane images, their corresponding nucleus stacks were processed by AceTree, which can be used to identify cell’s name and the lifespan of each nucleus. We defined volume consistency and ratio of lost nuclei to estimate the error of segmentations in four wide-type embryos with 200 frames in each, which are used for following spatiotemporal reference reconstruction. A index matrix **V**_*tc*|*t*∈[1,2,3,…200],*c*∈*C*_ was constructed such that “0” represents the existence of cell *c* at time *t*, otherwise the element is kept as *NaN* (invalid value), where *C* is the collection of cell names. We assembled cell’s volume of all segmentations of one wide-type embryo into the matrix **V**. Then for each cell *c*, the volume consistency *ρ*_*c*_ is defined as the ratio of the standard deviation and mean applied to volume series **V**_*t,c*|*t*=1,2,…,200_. Note that all invalid values *NaN* were not taken into consideration. The start time point of cell *c* is also labelled as *t*_*c*_ in order to discriminate the error at different developmental stages. Larger *ρ*_*c*_ means the segmentation of cell *c* has better temporal consistency, yielding higher performance. The distribution of consistency coefficients (*t*_*c*_, *ρ*_*c*_) of the four wide-type embryos is plotted in Fig. 1**B**. In these segmentations, most cells have relative small volume variation through the development, although temporal information are not applied to CShaper in the segmentation procedures intentionally. With the number of cells increasing over time, the cell volume becomes smaller and the signal-to-noise ratio decreases dramatically, which makes it more challenging to be segmented precisely. Simultaneously, because nuclei are not involved in the segmentation stage, nucleus information processed by AceTree can be used to justify the accuracy. In Fig. 1**C**, we show the ratio of lost nucleus (see **Supplementary note 1**) at different time points. Few cells before 200-cells stage are lost in the four embryos. With the number of cells increasing and density of fluorescence shrinking, the number of cells that are lost becomes larger when entering the eighth round of cell divisions. However the lost cells in the entire embryos only occupy a small proportion (around 11.9% at 350-cells). There-fore, we can safely conclude that CShaper has superior ability on segmenting extensive time-lapse embryo images.

### Establishment of spatio-temporal morphogenesis reference

Using experimental methods and quality-control standards described before^6,60^, 4 wild-type embryos within 350-cell stage were used to construct the morphogenesis reference of early *C. elegans* embryo. All the 17 embryos (4 embryos with both nucleus and membrane markers and the other 13 embryos with only nucleus marker) were first linearly normalized to minimize their positional variation according to a proposed computational pipeline^60^; secondly, translation in *yz* plane and rotation around *x* axis were successively performed on the 4 embryos with membrane marker, to ensure the compressed contact faces were parallel to *xz* plane; thirdly, translation in *xz* plane and rotation around *y* axis were successively performed on the 4 embryos, so that their projection to *xz* plane could be embedded by a centralized ellipse with minimum area; after that, the 4 embryos were rescaled to the same length in *x, y, z* directions; at last, the other 13 embryos were normalized to the nucleus-position averages of the 4 embryos using methods proposed previously^60^ (**Supplementary** Fig. S2).

### Morphological developmental properties at single-cell level

Using the cell-segmentation algorithm designed above, a total of 226 cells (≈ 70%) among AB4-128, MS1-MS16, E1-E8, C1-C8, D1-D4, P3 and P4 were recorded with complete lifespans and segmented without any error in all the 4 wild-type embryos, generating 4-dimensional morphological dynamic information at single-cell level with high confidence, such as cell shape (e.g. cell volume, cell surface area, topology) and cell-cell contact (e.g. contact duration, contact area, neighbour relationship) (Fig. 2, **Supplementary** Fig. S3)(Table. S2).

**Figure 2:**
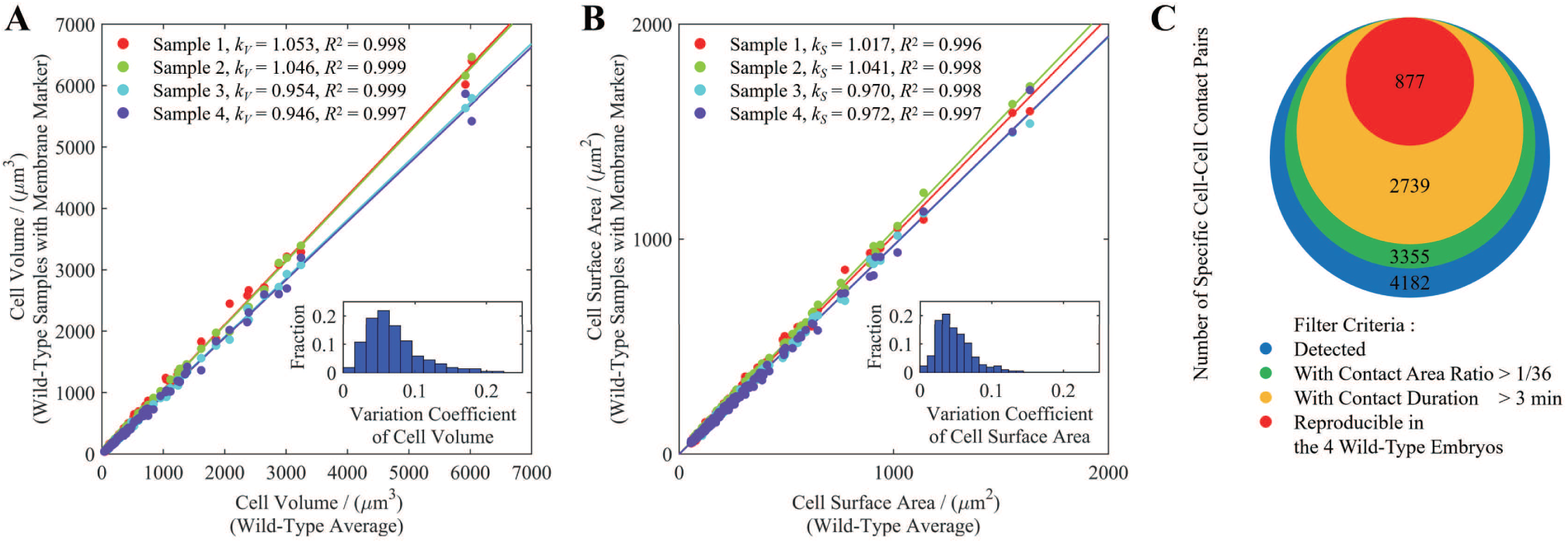
Reproducibility, precision, validity of cell volume, cell surface area, and cell-cell contact. All the cells involved are completely recorded and segmented without any error during their lifespans (Table 1). (**A**) Highly reproducible volume of cells in the 4 wild-type embryos. Inset, variation coefficients of cell volume. (**B**) Highly reproducible surface area of cells in the 4 wild-type embryos. Inset, variation coefficients of cell surface area. (**C**) Selection filter of sufficient, continuous and reproducible cell-cell contact pairs according to a set of arbitrary criteria.

To test the data quality and further refine the information useful to biological study, here we focus on three low-dimensional but valuable developmental properties: cell volume, cell surface area and cell-cell contact. Cell volume and cell surface area, which usually get involved in cell-cycle control and cell-fate specification^13–18^, were firstly investigated and found to have high reproducibility among individuals (Fig. 2 **A**,**B**). Cell-fate induction (e.g. Wnt signaling^19,20^ and Notch signaling^10,61^) during embryo development intimately depends on continuous, sufficient and direct physical contact between specific cells for interaction between receptors and ligands and consequent effective signaling transmission. Based on the contact relationship acquired from raw experimental images and automatic segmentation, we filtered the most reliable and valid contact between two specific cells by adding three empirical criteria^10^, **1)** with contact area larger than 1/36 of at least one cell’s surface area (sufficiency, *S*_**contact**_*/S*_**surface**_ > 1*/*36); **2)** with contact duration no shorter than 3 minutes, i.e. consecutive two time points (continuity, *T*_**contact**_ ≥ 3 min); **3)** be reproducible in all the 4 wild-type embryos (reproducibility, *N*_*replicate*_ = *N*_*embryo*_). At last, totally 877 contact pairs of two specific cells are identified (Fig. 2**C**). Cells with missing information are listed in **Supplementary** Table. S3. Please note that the criteria are set arbitrarily and can be readjusted for different research purposes.

**Table 1:** Verification of our newly proposed morphogenesis reference using experimental results from references.

| Cell-Cell Contact Pair |  |  | Validity<br>&<br>Reproducibility | Contact<br>Duration<br>(min) | Relative<br>Contact Area<br>Surface (%) | Relative<br>Contact Area<br>Surface (%) | Remark | Ref. |
| --- | --- | --- | --- | --- | --- | --- | --- | --- |
| P2 | → | EMS | 4/4 | 7.04 | 9.61±2.30 | 12.31±2.15 | Wnt Signaling | Thorpe et al<sup>19</sup> |
| MS | → | ABal | 4/4 | 7.04 | 15.70±2.60 | 15.03±2.66 | Latrophilin Signaling | Langenhan et al<sup>63</sup> |
| C | → | ABar | 3/4# | 7.04 | 2.37±2.57 | 2.60±2.79 | Wnt/β-catenin Signaling | Walston et al<sup>20</sup> |
| P2 | → | ABp | 4/4 | 5.63 | 13.38±1.99 | 20.47±2.01 | Notch Signaling (1<sup>st</sup>) | Priess et al<sup>61</sup> |
| MS | → | ABalp | /* | / | / | / | Notch Signaling (2<sup>nd</sup>) | Chen et al<sup>10</sup>, Priess et al<sup>61</sup> |
| MS | → | ABara | 4/4 | 5.63 | 9.28±1.19 | 12.98±2.14 | Notch Signaling (2<sup>nd</sup>) | Chen et al<sup>10</sup>, Priess et al<sup>61</sup> |
| ABalapa | → | ABplaaa | 4/4 | 5.63 | 14.22±3.86 | 11.13±3.26 | Notch Signaling (3<sup>rd</sup>) | Chen et al<sup>10</sup>, Priess et al<sup>61</sup> |
| ABalapp | → | ABplaaa | 4/4 | 5.63 | 13.06±3.91 | 10.66±2.65 | Notch Signaling (3<sup>rd</sup>) | Chen et al<sup>10</sup>, Priess et al<sup>61</sup> |
| MSapp | → | ABplpapp | 4/4 | 5.63 | 12.58±3.69 | 13.92±3.86 | Notch Signaling (4<sup>th</sup>) | Chen et al<sup>10</sup>, Priess et al<sup>61</sup> |
| MSapppp | → | ABplpppp | 4/4 | 5.63 | 7.58±0.21 | 9.19±2.61 | Notch Signaling (5<sup>th</sup>) | Chen et al<sup>10</sup>, Priess et al<sup>61</sup> |

## Discussion

Cell morphology is critical and useful developmental property involved with different biological process. In this paper, a complete pipeline CShaper for analyzing spatio-temporal morphological features of *C. elegans* embryo at single-cell level is proposed. The CShaper benefits from the well-defined distance learning task. By learning to capture multiple discrete distances, DMapNet extracts the membrane mask by considering shape information, instead of just intensity features. Remarkable performance is examined by both intrinsic geometric constrains and previous notable discoveries. Based on these accurate segmentations, we merged the embryos and quantitatively generated a spatio-temporal morphogenesis reference for 4-to 350-cell stage of *C. elegans* embryogenesis, including single-cell properties such as cell shape (e.g. cell volume, cell surface area, topology), cell-cell contact (e.g. contact duration, contact area, neighbour relationship), cell positional variability, etc. In all, 226 cells with complete lifespans and dynamic morphology trajectory were produced. Furthermore, a total of 877 contact pairs between two specific cells were identified with high reproducibility, continuous contact duration and sufficient contact area, which could be a solid foundation for searching potential signaling pathways between cells. Our work provides a quantitative and statistical framework for *C. elegans* morphogenesis, which is a powerful resource to push forward multiple biological research fields like signaling transmission, fate specification and asymmetric segregation. Next, we discuss the coincidence of CShaper with three well-known experimental discoveries and inspect its valuable applications on different datasets.

### Verifications on previous studies

To further validate the *C. elegans* morphogenesis reference with single-cell developmental properties (e.g. cell volume, cell position, cell-cell contact), here we use our new quantitative data to repeat and verify three separate sets of experimental conclusions about *C. elegans* embryonic development^10,13,61–63^.

Firstly, Arata et al. found the power law relationship between cell cycle duration and cell volume in the early *C. elegans* development, that is, AB and MS cells adopt the same power exponent (≈-0.27) while C and P cells adopt another different power exponent (≈-0.41)^13^. Under log-log scale coordinates system, we also performed linear fitting between cell cycle duration and cell volume and found that the two exponents obtained (AB and MS, −0.2990; C and P, −0.4244) are very close to the values proposed before (Fig. 3**A, B**).

**Figure 3:**
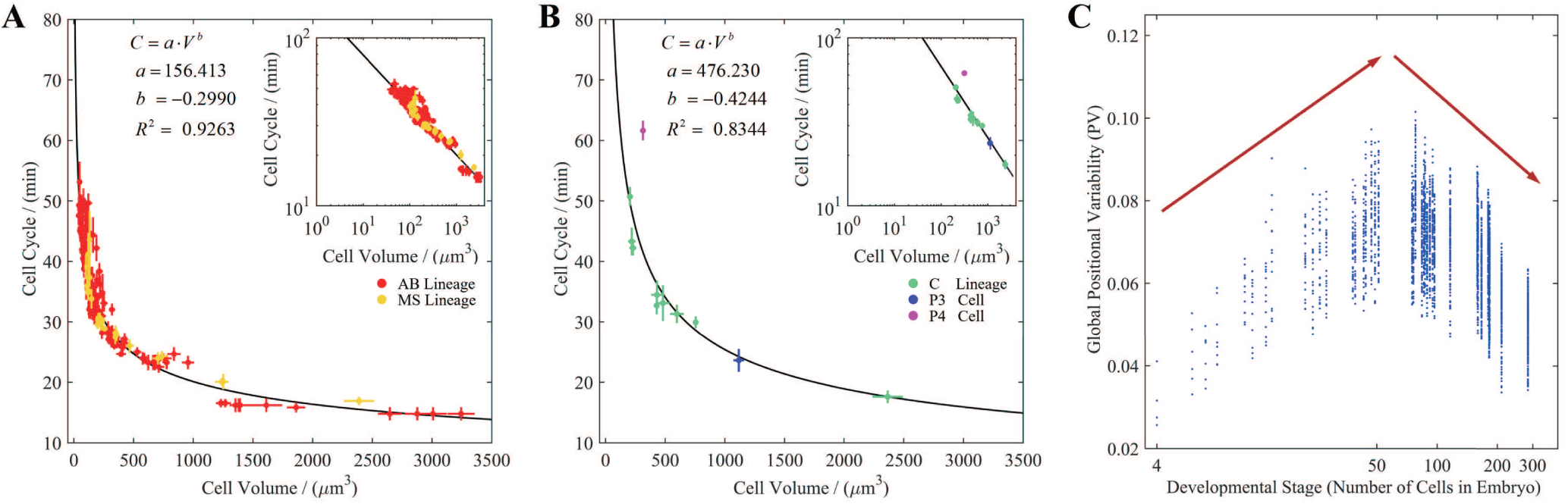
Verification of our newly proposed morphogenesis reference using experimental results from References^13,62^. (**A**) Power law relationship between cell cycle duration and cell volume in AB and MS cells, with power exponent ≈-0.2990. Inset, illustration in log-log scale coordinates system. (**B**) Power law relationship between cell cycle duration and cell volume in C and P cells, with power exponent ≈-0.4244. Inset, illustration in log-log scale coordinates system. (**C**) “Low-high-low” dynamic pattern of cell positional variability during 4- to 350-cell stage.

Secondly, Li et al. uncovered the “low-high-low” dynamic pattern of cell positional variability during 4-to 350-cell stage of *C. elegans* embryogenesis^62^. Using the same evaluation method on positional variability (RMSD) proposed before, we used our normalized cell positions (nucleus positions) to calculate their spatial variation and reconstructed a similar curve with a turning point occurring when cell number reaches around ninety (Fig. 3**C**).

Thirdly, several intercellular signaling based on accurate cell-cell contact have been identified to play important roles in asymmetric segregation, spindle formation and cell-fate induction^10,19,20,61,63^. Here, we compared the known cell-cell signaling pairs with our cell-cell contact dataset (877 pairs in total). The majority of known contact pairs past through our filter criteria with continuous contact duration as well as sufficient contact area, except C → ABar and MS → ABalp (Table. 1). The contact between C and ABar can be found in all the 4 wild-type embryos with at least two consecutive time point (≈3 minutes), however, the relative contact area is smaller than the arbitrary filter criterion (*S*_**contact**_*/S*_**surface**_ > 1*/*36) in one of the embryos (Fig. 2**C**), revealing that the threshold for identifying valid cell-cell contact should be reestimated and readjusted on the basis of the actual biological scenes (e.g. surface density of ligands or receptors), nevertheless, the original contact information from image segmentation is objective and can be used for different research purposes. For the other contact pair MS → ABalp, one of the cells ABalp are missing (i.e. fail to be segmented) due to its location near the top of embryo and consequent dim fluorescent signal.

### Application on different datasets

To evaluate the performance of CShaper on different kinds of images, the plant tissue images used by Fernandez et al^64^ were segmented with our method. In the work, Arabidopsis thaliana stem cells were processed with MARS^65^, which provides reasonable discrimination on inner parts of the tissue by fusing multi-angle acquisitions. Different from MARS, CShaper processes the stem in a more challenging way, segmenting single-direction membrane stack without the fusion stage. Because of the large shape difference between animal and plant cells, we retrained DMapNet with two segmentation results from the MARS. Negative segmentations were filtered when its volume deviates the average level too much. The comparison result is listed in **Supplementary** Fig. S4. Both MARS and CShaper show promising segmentations of cells at shallow layers. However, CShaper shows stronger results on inner parts with extremely low intensity. Although DMapNet was trained with defective reference as shown in **Supplementary** Fig. S4**B, C** and **D**, CShaper escaped these irregular errors during the inference stage.

### Constrains of CShaper

CShaper provides new insights into the study of *C. elegans* embryogenesis at single-cell level, both in spatial and temporal aspects. First, to promote the accuracy of automatic *C. elegans* embryo segmentation, especially at later developmental stage, some constrains of CShaper need to be emphasized here. First, as the cell shape changes with time continuously, the temporal features between consecutive frames are supposed to elevate the segmentation performance. LSTM, originally designed for natural language process (NLP), is an obvious candidate to capture temporal features across time^26^. However, CShaper doesn’t adopt LSTM-based model, such as convLSTM^66^, because of the considerable computational resources involved in 3D convolution. We also find that the segmentation errors of CShaper concentrate at the top of the embryo, where the membrane signal intensity decreases critically because of the laser attenuation. In the framework of CShaper, potential strategies could be used to normalize the quality of top half embryo based on the button one. For example, Generative Adversarial Networks (GAN) can be employed to transform low-quality images into the target with higher resolution^40,45^.

## Methods

CShaper consists of three phases. The first step is to extract the membrane mask by deep learning based DMapNet. After that, Delaunay triangulation is used to cluster local minimum for the followed watershed segmentation. The negative segmentations and potential gaps among cells are filtered with the nucleus channel. The final cell shape lineage is constructed based on the series information provided by the nucleus lineage. The framework of CShaper is shown in Fig. 4.

**Figure 4:**
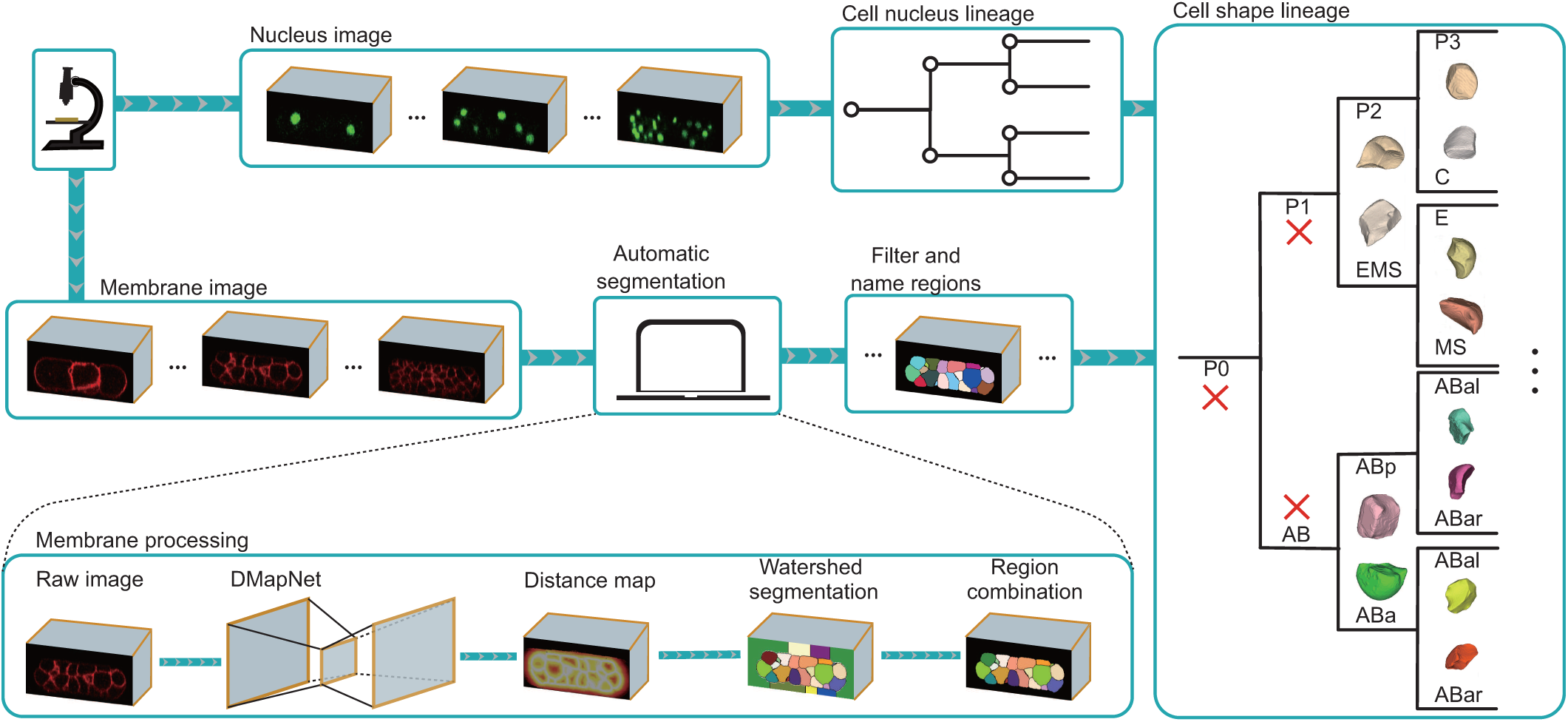
The framework of CShaper. Serial nucleus and membrane stacks are imaged simultaneously. For membrane image, CShaper is applied to segment the embryo at single-cell level automatically. Nucleus stack is processed by AceTree, which tracks nucleus through the entire development process. Finally, CShaper embedded cellular shape into the nucleus lineage.

### Distance Constrained Learning

The noise and physical imaging limitation degrade the quality of automatic segmentations. This problem prevails in embryo segmentation since the membrane enclosing a cell can be hardly imaged completely. DMapNet adopts a distance constrained learning to address the problem in segmenting noisy embryo images. By implicitly learning the distance map, the DMapNet is able to discriminate weak or even lost membrane signal.

With the membrane labelled as front label 1, the binary membrane annotation *M ^B^* was prepared as discrete distance map *M ^D^* with

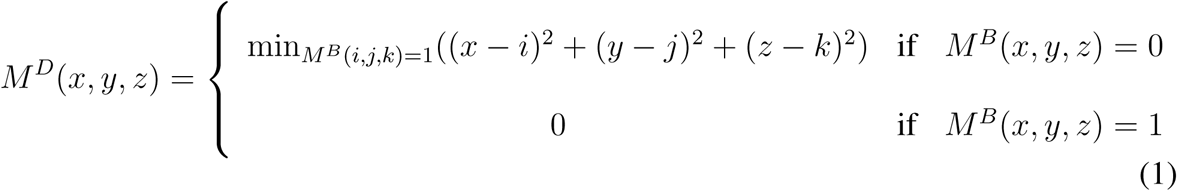

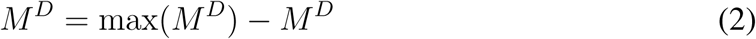

In Eq. (2), we reversed the distance map to keep it monotonically decreasing from the membrane to the background. To relieve the burden on learning the distance where pixel was too away from the membrane to have recognizable signal, *M ^D^* was clipped to prevent long-range spatial dependencies. Then *M ^D^* was further discretized nonlinearly into *M* ^*dmap*^, the learning target, with smaller intervals around the membrane mask. The cross-entropy loss was used to evaluate the learning progress. However, different from traditional multi-classification, DMapNet should ideally provide a class distribution such that the predicted class closer to the real class has higher probability than that is further away. For example, the penalty of predicting *l* = 1 as *l* = 2 should be smaller than that of the prediction *l* = 15. Therefore, the cross-entropy loss is weighted by the class distance as

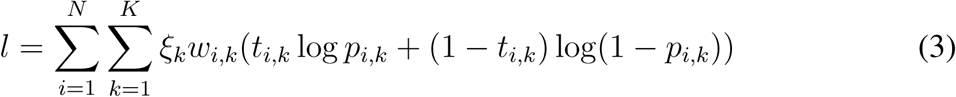

where *ξ*_*k*_ is the weight for each interval, *w*_*i,k*_ is the class distance weight for each pixel *i, t*_*i,k*_ is the *k*-th element of the one-hot target vector at pixel *i*, and *p*_*i,k*_ is the *k*-th channel of the network output at pixel *i. N* and *K* are the number of pixels and distance intervals, respectively. The *K*-th interval denotes the mask at the center of the membrane, while 0-th class represents the background far away from the membrane. Larger weight *w*_*k*_ on classes closer to the membrane helps the network put more attention on cell boundary. The class distance weight *w*_*i,k*_ is calculated as

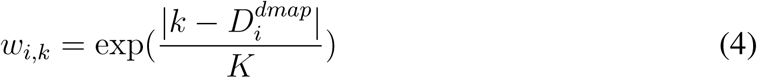

where 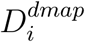 is the real class at pixel *i*.

### Network structure

The structure of DMapNet is shown in **Supplementary** Fig. S5. Taking the fully convolutional network as the backbone, DMapNet is constructed by considering some specific problems in fluorescent images. In order to reduce 3D computational complexity, DMapNet utilizes inter-slice information by using 3 × 3 × 1 and 1 × 1 × 3 convolution successively. The dilation convolution, from the output of inner-slice convolution to its input, is added to enlarge the receptive filed. The residual block is composed of two convolutional layers^67^. Parametric Rectified Linear Units (PReLU)^68^ are used as the activation layer. To help the higher layers retain the information from pixel, the input is scaled down and concatenated with the feature map, which also helps to train the network on cells in different sizes^51^. This is essential for segmenting cells when annotations corresponding to late development stage are not reliable. Feature maps at lower resolution are upsampled and concatenated together to generate the probability map. The membrane image volume is split into multiple overlapped slice series *I*_*D*×*W* ×*H*_, which are processed by DmapNet individually. The final prediction of whole volume 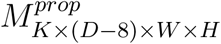 is achieved by packing these predictions sequentially as discussed in Distance Constrained Learning. Then the discrete distance map can be derived by *M* ^*pred*^(*z, x, y*) = max_*c*=[1,…*K*]_ *M* ^*prob*^(*c, z, x, y*).

### Automatic seeds clustering based on Delaunay triangulation

Till now, we have just obtained the discrete distance map *M* ^*pred*^ from the DMapNet, as shown in Fig. 4. Watershed segmentation is well suited for separating individual cells from the map, where black cell interiors are surrounded by bright boundaries. The learned map *M* ^*pred*^, including multiple discrete distance intervals, approximates the distance transformation on the latent binary membrane mask. With the holistic information embedded in the ordered intervals, cell boundaries can be extracted. However, watershed algorithm cannot be applied on *M* ^*pred*^ directly because of redundant local minima and low distance resolution. In this part, Delaunay triangulation is employed to detect seeds for watershed segmentation automatically.

First, by selecting the *K*-th interval as the foreground mask 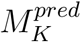, a reversed EDT was applied to 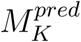, yielding *I*^*edt*^. All local H-minima in *I*^*edt*^ are noted as 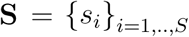, where *S* is number of local minima. In order to filter *s*_*i*_s that belong to the same cell or background, a weighted graph **G** is constructed. Edges **E** = {**E**_1_, **E**_2_} in **G** come from two sources: one is the Delaunay triangulation on **S**, noted as **E**_1_; another one is the edges **E**_2_ among all local minima locating on the boundary of the volume. Weight of the edge **e**_*ij*_ is defined as

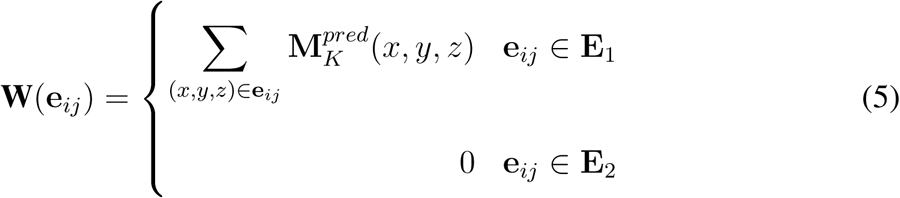

where (*x, y, z*) ∈ **e**_*ij*_ represents all points on the edge **e**_*ij*_. The edge is moved from **E** if its weight is greater than the OTSU^69^ threshold on **W**. Finally, vertexes **M** were clustered based on their connectivity. All *s*_*i*_s in one cluster are regarded as one seed in the watershed segmentation stage. In the seeding procedure, a seed could possibly locate in a hollow gap between cells, giving rise to fake cell regions. Thus, nucleus stack was used to modify these errors. Owing to the impressive performance of DMapNet and automatic seeding, only intensity normalization was needed for nucleus image. Regions were set as background when the cumulated intensity in the region is smaller than a threshold.

## Acknowledgements

The strains were provided by the *C. elegans* Genetic Center (CGC), which is funded by National Institutes of Health, Office of Research Infrastructure Programs, Grant P40 OD010440. This work was supported by the Hong Kong Research Grants Council (HKBU12100118, HKBU12100917, HKBU12123716), the HKBU Interdisciplinary Research Cluster Fund, the Ministry of Science and Technology of China (2015CB910300), the National Natural Science Foundation of China (91430217), Hong Kong Research Grants Council (Project C1007-15G) and City University of Hong Kong (Project 9610460).

## Author Contributions

H.Y., Z.Z., C.T. conceived and coordinated the study. J.C. designed the cell-segmentation algorithm; G.G. analysed the segmentation results. M.W., L.C. performed imaging and embryo curation. Z.Z. provided reagents and experimental methods. J.C., G.G. wrote the manuscript; H.Y., Z.Z., C.T. revised the manuscript. All the authors reviewed the results and approved the final version of the manuscript.

## Competing Interests

The authors declare that they have no competing financial interests.

## Supplementary

**Figure S1:**
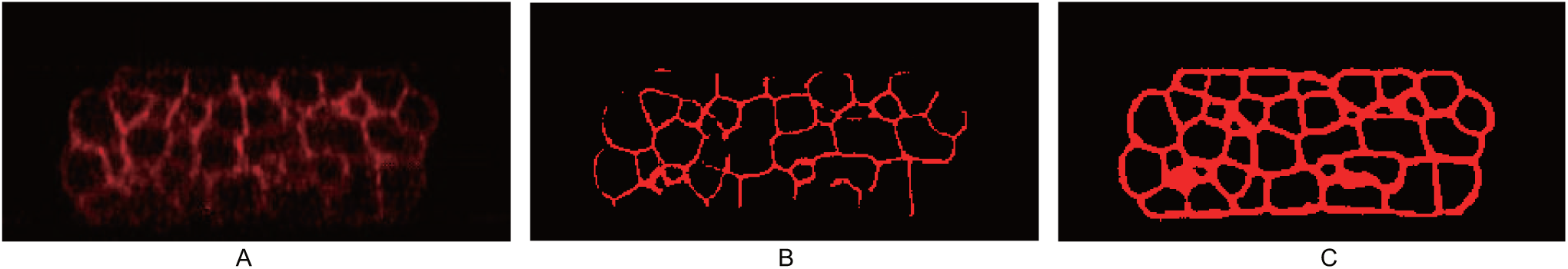
Comparison of CShaper and naive binary segmentations. (**A**) Raw image. (**B**) and (**C**) are segmentations of naive binary and CShaper, respectively.

**Figure S2:**
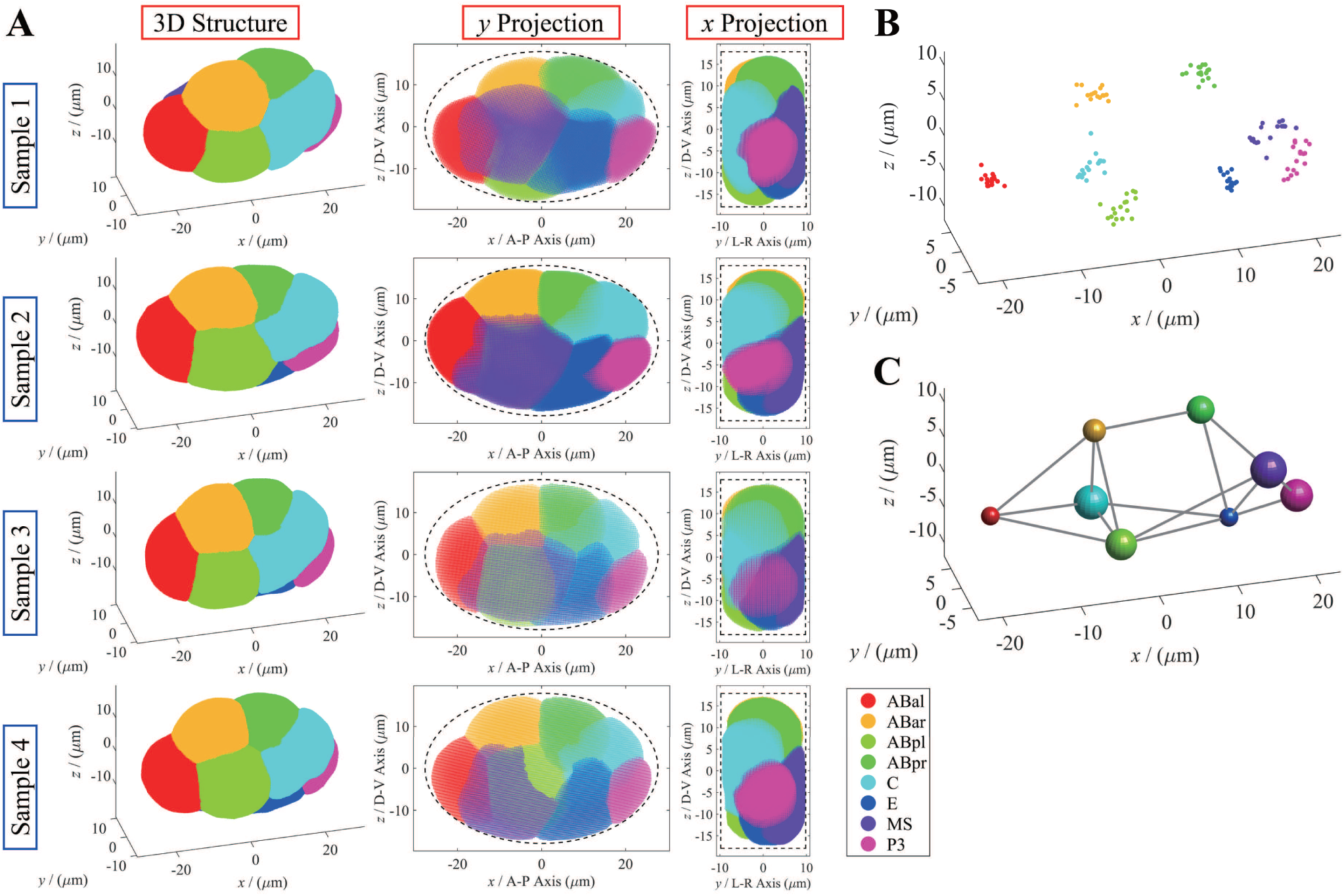
Spatio-temporal reference of wild-type *C. elegans* embryonic morphogenesis. Take the 8-cell stage for example and illustration. *x, y, z* axes represent anterior-posterior (A-P), left-right (L-R), dorsal-ventral (D-V) axes respectively. Each color represents one specific cell identity, noted in legend. (**A**) 3D structure, *y* projection and *x* projection of cell morphology in 4 wild-type embryo samples. (**B**) Distribution of nucleus position in 17 wild-type embryo samples. Each point represents a cell’s nucleus position in one embryo sample. (**C**) Spatial deviation and cell-cell contact mapping. Radius of sphere represents spatial deviation Δ*r*_STD_ defined by 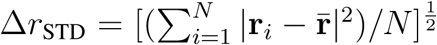; gray lines represent reproducible and effective contact between cells, under arbitrary filter criteria (*S*_contact_*/S*_surface_ > 1*/*36; *T*_contact_ ≥ 3min; *N*_replicate_ = *N*_embryo_).

**Figure S3:**
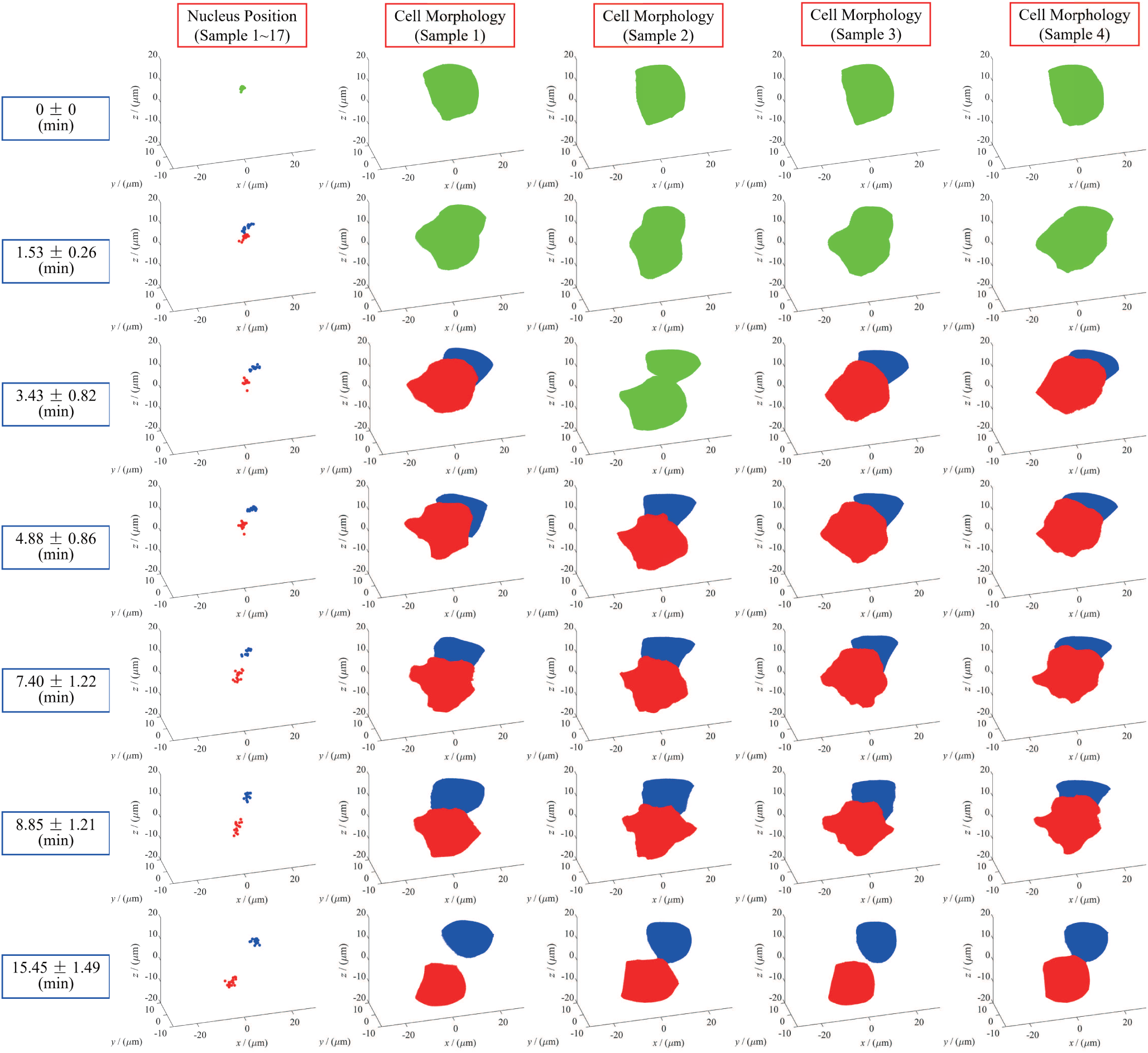
4-dimensional morphological evolution during *C. elegans* embryogenesis. Take ABp and its daughters ABpl, ABpr as example and illustration. *x, y, z* axes represent anterior-posterior (A-P), left-right (L-R), dorsal-ventral (D-V) axes respectively. Green, ABa; red, ABpl; blue, ABpr. Evolution dynamics is shown in rows which represent different developmental timing noted on left (Tabel. S2). The first column is nucleus-position distribution from 17 wild-type embryo samples. The second to fifth columns are reconstructed cell morphology from the 4 wild-type embryo samples.

**Figure S4:**
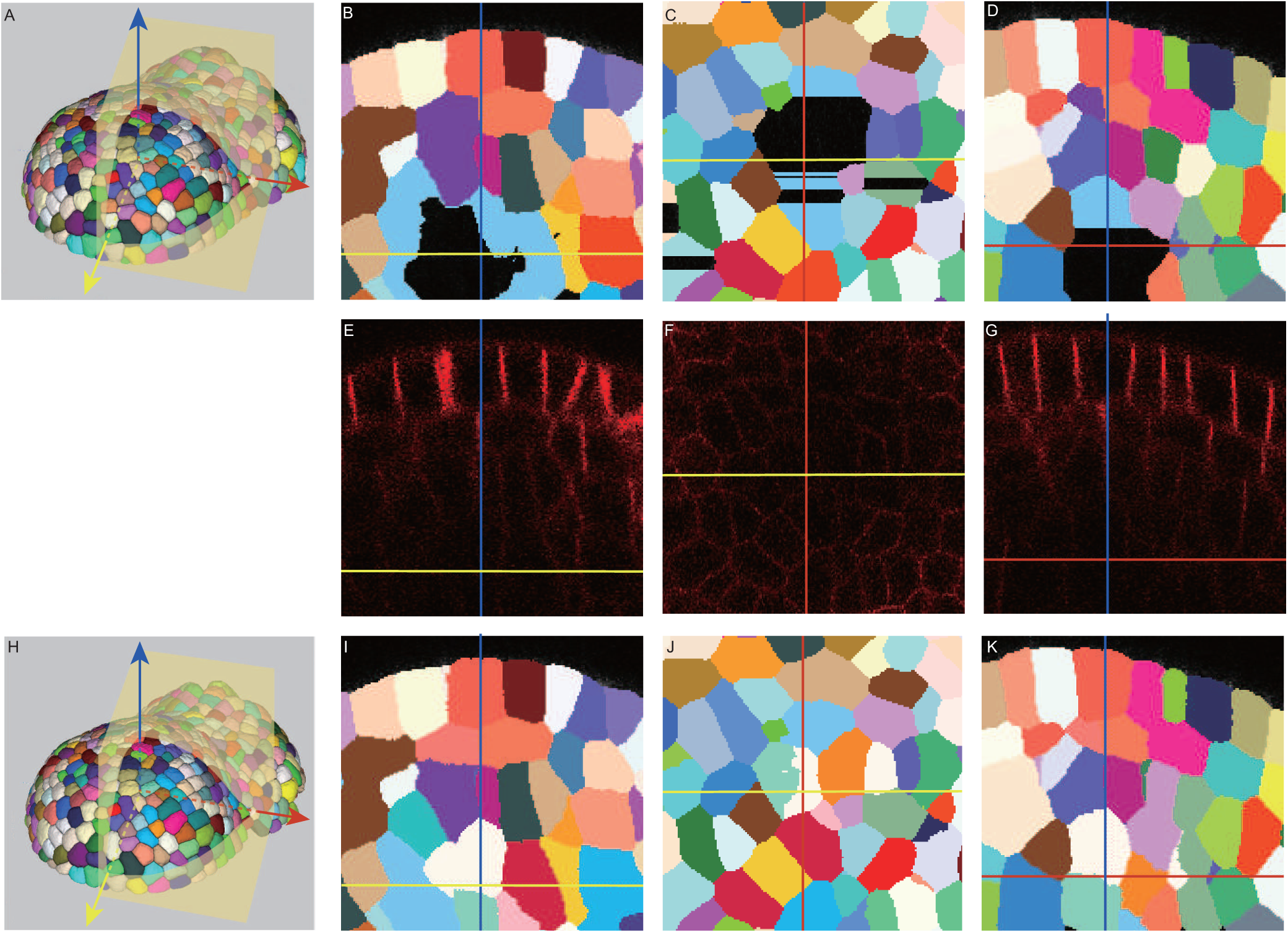
The application of CShaper on plant tissue. (**A, H**) are segmentations of MARS and CShaper shown in 3D. (**B-D**, (**E-G**) and (**I-K**) are three orthogonal sections of MARS’s segmentation, raw image and CShaper’s segmentation, respectively.

**Figure S5:**
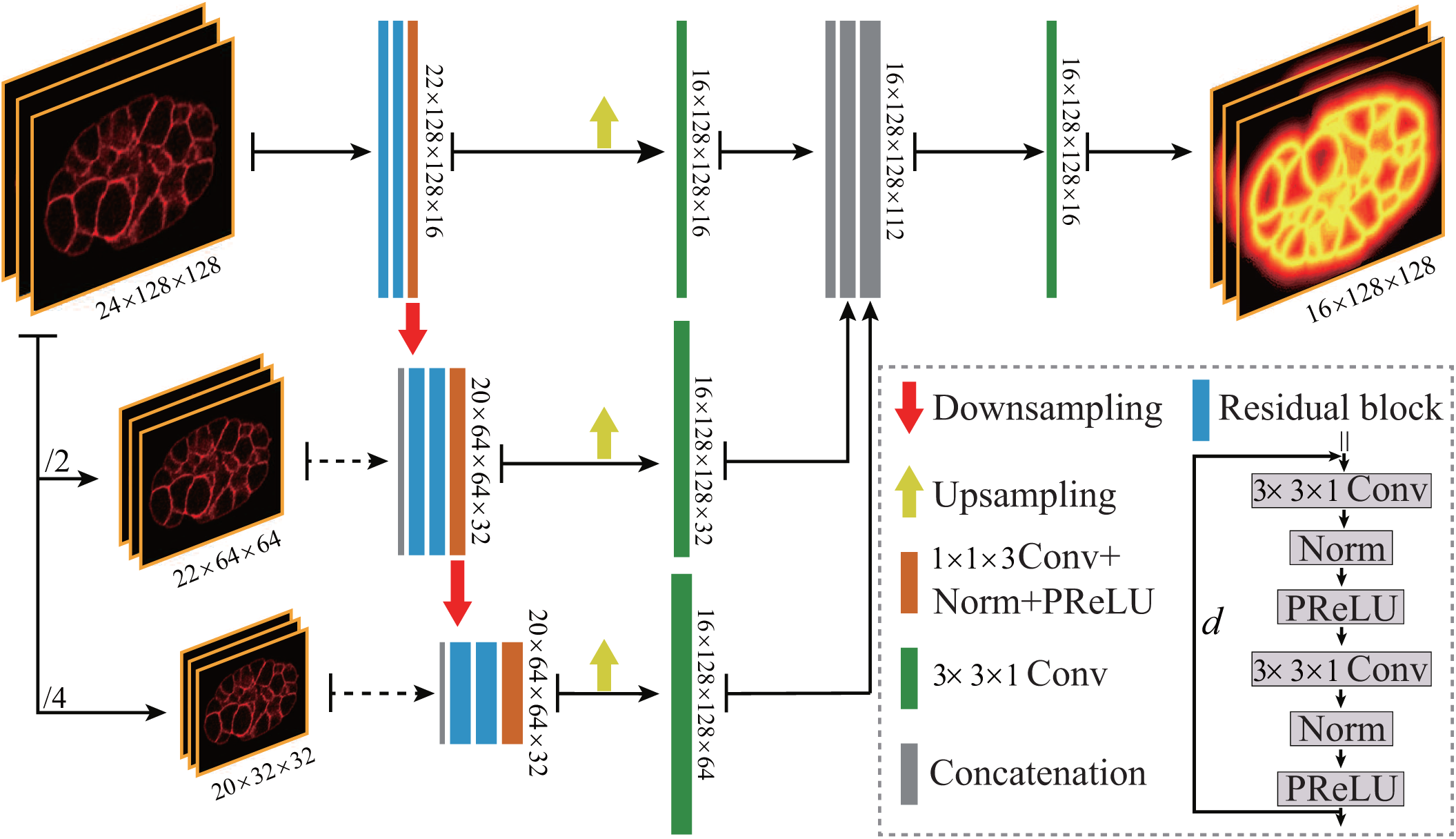
Network structure of DMapNet. Multiple neighboring slices are processed by three residual blocks at three different levels consecutively. Feature maps at high levels are resized into the same size as the input image with linear interpolation. In order to remedy the lost information, raw images are downsampled and concatenated into the feature maps. The complete distance map of one volume is achieved by combining multiple predictions.

**Table S1:**
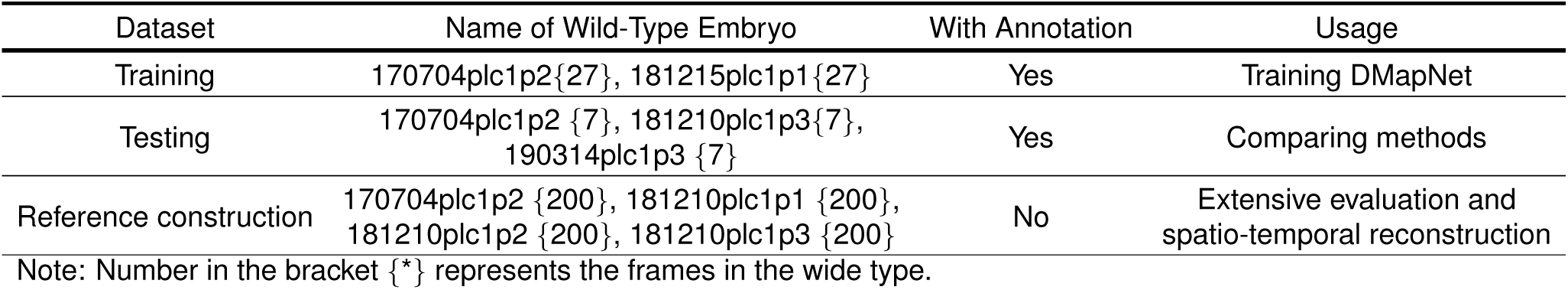
Datasets (with membrane marker) description

| Dataset | Name of Wild-Type Embryo | With Annotation | Usage |
| --- | --- | --- | --- |
| Training | 170704plc1p2{27}, 181215plc1p1{27} | Yes | Training DMapNet |
| Testing | 170704plc1p2 {7}, 181210plc1p3{7},<br>190314plc1p3 {7} | Yes | Comparing methods |
| Reference construction | 170704plc1p2 {200}, 181210plc1p1 {200},<br>181210plc1p2 {200}, 181210plc1p3 {200} | No | Extensive evaluation and<br>spatio-temporal reconstruction |

Note: Number in the bracket {\*} represents the frames in the wide type.

**Table S2:** Time segmentation on developmental process from 4-to 350-cell stage.

| Division Event | First Moment<br>(min) | Last Moment<br>(min) | Division Event | First Moment<br>(min) | Last Moment<br>(min) |
| --- | --- | --- | --- | --- | --- |
| AB2 | / | 0.00±0.00 {01} | E1 | 4.88±0.86 {04} | 22.29±1.82 {10} |
| AB4 | 1.53±0.26 {02} | 15.45±1.49 {07} | E2 | 23.74±1.87 {12} | 62.21±4.53 {25} |
| AB8 | 17.67±1.30 {08} | 32.25±2.83 {15} | E4 | 65.69±5.01 {26} | 106.92±5.34 {39} |
| AB16 | 35.21±3.22 {18} | 56.47±3.80 {23} | E8 | 113.14±5.66 {41} | 175.11±6.86 {52} |
| AB32 | 60.92±4.11 {24} | 84.84±5.15 {32} | E16 | 191.30±6.79 {54} | / |
| AB64 | 92.25±5.15 {34} | 118.51±5.60 {44} | C1 | 8.85±1.21 {06} | 25.92±2.60 {13} |
| AB128 | 134.41±6.13 {46} | 159.97±4.64 {50} | C2 | 27.37±2.61 {14} | 49.85±3.47 {21} |
| AB256 | 190.98±4.70 {53} | / | C4 | 51.91±3.65 {22} | 81.15±4.85 {31} |
| EMS | / | 3.43±0.82 {03} | C8 | 88.20±5.55 {33} | 115.85±6.12 {43} |
| P2 | / | 7.40±1.22 {05} | C16 | 143.16±7.27 {47} | / |
| MS1 | 4.88±0.86 {04} | 21.18±1.71 {09} | D1 | 34.38±2.90 {16} | 68.21±5.97 {28} |
| MS2 | 22.63±1.76 {11} | 42.00±3.12 {19} | D2 | 69.66±5.98 {29} | 112.53±6.83 {40} |
| MS4 | 43.96±3.43 {20} | 67.75±4.55 {76} | D4 | 115.77±7.34 {42} | 155.45±7.15 {49} |
| MS8 | 72.20±4.91 {30} | 97.58±5.76 {35} | D8 | 163.81±7.13 {51} | / |
| MS16 | 104.74±6.14 {38} | 130.54±6.05 {45} | P3 | 8.85±1.21 {06} | 32.93±2.88 {16} |
| MS32 | 153.27±6.10 {48} | / | P4 | 34.38±2.90 {17} | 99.90±6.89 {36} |
|  |  |  | Z2/Z3 | 101.35±6.90 {37} | / |

Note: Number in the bracket{\*} represents the order of the 54 division events (frames).

**Table S3:**
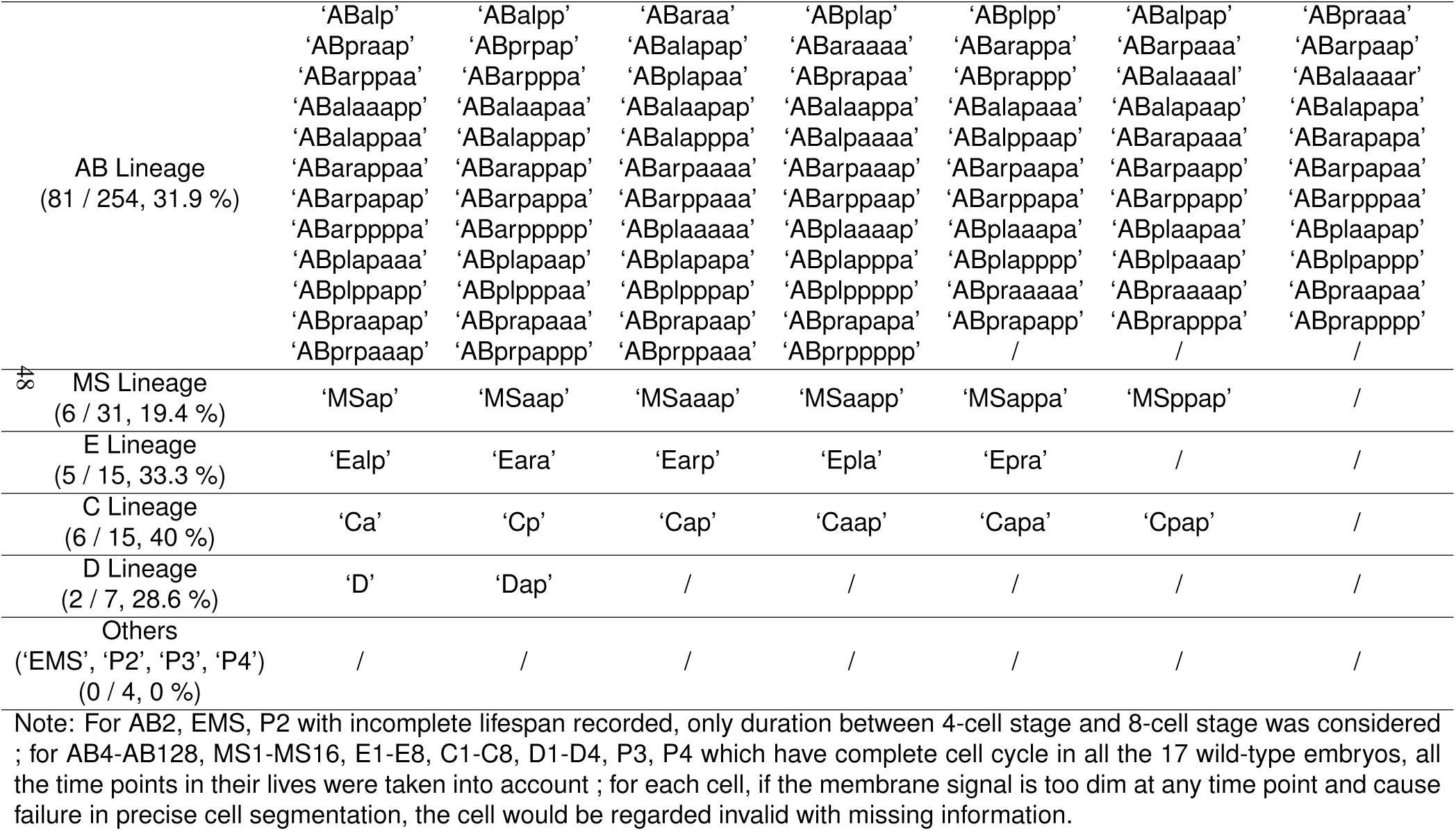
Cells with missing information during 4-to 350-cell stage in the 4 embryos expressing membrane marker (30.7%).

**Note 1**: In CShaper, there are three discriminative situations where nucleus derived from AceTree cannot be found:

1. The boundary between two cells (not sisters) is too week to be extracted by DMapNet, therefore, two cells are segmented as one cell during the watershed transformation;
2. Membrane signal is lost at boundary of the embryo, which leads to the leakage of the background into the embryo;
3. In CShaper, we determine the accomplishment of division stage by checking the signal intensity on the line between sister cells’ nuclei. However, when the intensity drops at the middle of lifespan, the sister cells may be combined as their mother cell, leading to missing cells.

We exclude these mistakes by combining the nucleus lineage from AceTree and segmentations from CShaper.

